# Bixafen, a succinate dehydrogenase inhibitor fungicide, causes microcephaly and motor neuron axon defects during development

**DOI:** 10.1101/2020.08.15.252254

**Authors:** Alexandre Brenet, Rahma Hassan-Abdi, Nadia Soussi-Yanicostas

## Abstract

Succinate dehydrogenase inhibitors (SDHIs), the most widely used fungicides in agriculture today, act by blocking succinate dehydrogenase (SDH), an essential and evolutionarily conserved component of mitochondrial respiratory chain. Recent results showed that several SDHIs used as fungicides not only inhibit the SDH activity of target fungi but also block this activity in human cells in *in vitro* models, revealing a lack of specificity and thus a possible health risk for exposed organisms, including humans. Despite the frequent detection of SDHIs in the environment and on harvested products and their increasing use in modern agriculture, their potential toxic effects *in vivo*, especially on neurodevelopment, are still under-evaluated. Here we assessed the neurotoxicity of bixafen, one of the latest-generation SDHIs, which had never been tested during neurodevelopment. For this purpose, we used a well-known vertebrate model for toxicity testing, namely zebrafish transparent embryos, and live imaging using transgenic lines labelling the brain and spinal cord. Here we show that bixafen causes microcephaly and defects on motor neuron axon outgrowth and their branching during development. Our findings show that the central nervous system is highly sensitive to bixafen, thus demonstrating *in vivo* that bixafen is neurotoxic in vertebrates and causes neurodevelopmental defects. This work adds to our knowledge of the toxic effect of SDHIs on neurodevelopment and may help us take appropriate precautions to ensure protection against the neurotoxicity of these substances.

## Introduction

Bixafen, a methyl-pyrazole carboxamide, is a fungicide widely used on cereal and rapeseed crops. It was initially approved and marketed in 2011 by Bayer (Ravichandra 2018). Its high efficiency and rapid penetration resulted in a significant rise in usage in Europe and the US. Currently, nearly 30 different products are sold on the French market alone that contain bixafen as sole active substance or mixed with other fungicides. Bixafen belongs to the succinate dehydrogenase inhibitor (SDHI) family, the fungicides most widely used in agriculture to fight a broad range of fungal diseases (Lalève et al. 2014; Zhang et al. 2019). Bixafen, which is derived from carboxin and categorized as a latest-generation SDHIs, acts through inhibition of mitochondrial respiration chain complex II, also known as succinate dehydrogenase (SDH). However, the inhibitory effect of SDHIs is not restricted to fungi: they also block the mitochondrial respiration chain in many other species, including human cells (Bénit et al. 2019). Additionally, the latest-generation SDHIs have been shown to also inhibit mitochondrial respiration complex III (Bénit et al. 2019), suggesting more critical effects of these new fungicides. SDH activity is essential for proper mitochondrial respiration and metabolism, and defects in its activity impair the cellular metabolome and functions (Bénit et al. 2014). Many human diseases are related to SDH deficiency, including neurodegenerative disorders, such as Leigh syndrome, an early-onset neurodegenerative disease characterized by developmental delay, ataxia and seizures (Birch-Machin et al. 2000; Bourgeron 2015; Van Coster et al. 2003; Finsterer 2008; Horváth et al. 2006; Parfait et al. 2000), and cancers, such as familial paraganglioma syndrome, infantile leukoencephalopathy, and neuroblastoma (Ghezzi et al. 2009; Martin et al. 2007; Perry et al. 2006; Ricketts et al. 2009; Stratakis and Carney 2009; Timmers et al. 2009). The Norwegian scientific committee for food safety (VKM) has assessed the health and environmental risk of Aviator Xpro EC225, a fungicide containing bixafen, and concluded that the effects of bixafen in animals should be considered relevant for humans (VKM Report 2014). Also, M44, a bixafen metabolite, which has the potential to contaminate groundwater, causes abnormalities in the rabbit foetus, suggesting that similar effects in humans cannot be excluded (VKM Report 2015). However, despite the increasing use of SDHIs in modern agriculture and their frequent detection in the environment and on harvested products (Abad-Fuentes et al. 2015; Añasco, Koyama, and Uno 2010; Tanabe and Kawata 2009; Tsuda et al. 2009; Vu et al. 2016), the potential toxic effects of these substances, especially on neurodevelopment, have so far been poorly evaluated.

To fill this gap and assess the neurotoxicity of these latest-generation SDHI fungicides *in vivo,* we tested the potential toxic effect of bixafen *in vivo* on zebrafish embryos. We observed that embryos exposed to low or medium bixafen concentrations, LOAEL and LC30, developed a series of defects, including microcephaly and disorganized motor neuron axons and branches. This study demonstrated the toxicity of bixafen on CNS development and gives insight into the environmental risk of this SDHI on the nervous system in vertebrates.

## Results

### 1. Toxicity of bixafen *in vivo* in zebrafish embryos

To assess the potential toxicity of bixafen, an SDHI belonging to the pyrazole group of fungicides, zebrafish embryos were exposed to increasing concentrations of bixafen ranging from 0.1 μM to 4 μM, from 6 hours post-fertilization (hpf) onward up to 96 hpf (supplementary Figure 1A). As no differences in survival rates or embryo morphology could be detected between the 0.1% DMSO and fish water control groups (data not shown), embryos incubated in 0.1% DMSO were used as controls. Exposure of zebrafish embryos to 0.5 μM bixafen and above induced gross morphological defects and significantly increased mortality, compared with that seen in controls (supplementary Figure 1). Moreover, we observed a dose-dependent effect, with increasing concentrations of bixafen inducing increased mortality rates and increased phenotypic defects in surviving embryos (Supplementary Figure 1B_1_-K).

Results indicated that the median lethal concentration of bixafen was 2.7 μM for wild-type zebrafish embryos (AB line) after 96 hours of incubation, i.e. 96 hpf LC50, and closely similar values were observed with the two other zebrafish lines used in this study, the transgenic Tg[*Olig2:eGFP*] and Tg [*HuC:RFP*] (Supplementary Figure 1K).

To investigate the potential toxicity of bixafen, and particularly the effect of this SDHI on the morphology of the brain and spinal cord, we chose 0.2 μM, the lowest-observed-adverse-effect-level (LOAEL), and 0.5 μM, the 96 hpf LC30, two concentrations that induce mild and moderate phenotypic defects in 96 hpf embryos, respectively, except for small heart oedema in embryos exposed to 0.2 μM bixafen (Figure 1C), and large heart oedema and curved tail in embryos exposed to 0.5 μM bixafen (Figure 1D). We next carefully examined the morphological parameters of embryos exposed to 0.2 and 0.5 μM bixafen, namely body length, head size, eye diameter and eye-otolith distance. Results indicated that following exposure to either 0.2 or 0.5 μM bixafen, surviving embryos show reduced body length and eye diameter, and decreased head size and eye-otolith distance, compared with that observed in age-matched controls (Figure 1E-I).

**Figure 1:**
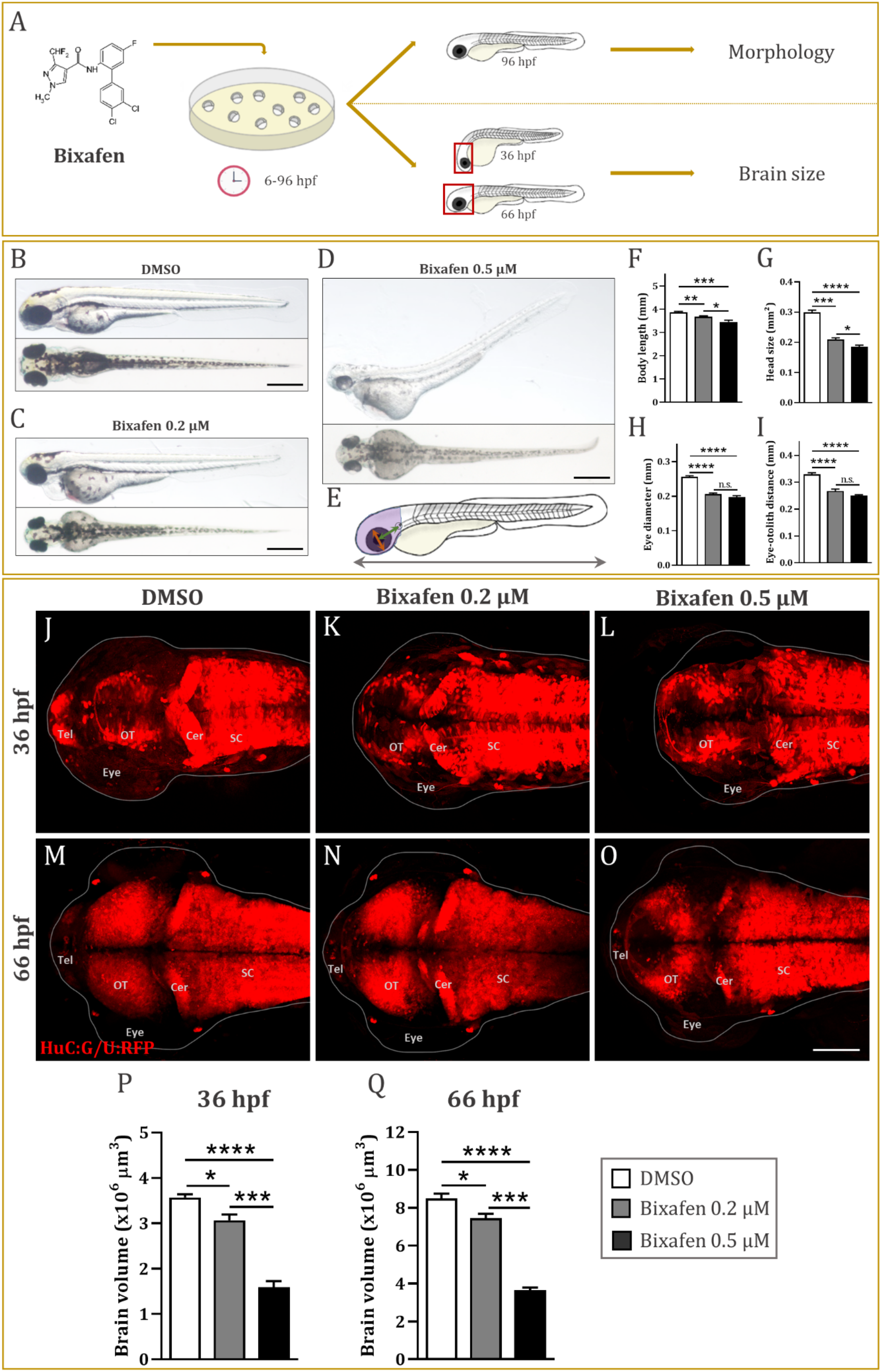
Bixafen exposure causes microcephaly. (**A**) Experimental setup used to analyze the body morphological parameters and brain size of zebrafish embryos exposed to 0.2 or 0.5 μM bixafen, from 6 hpf onward up to 36 or 66 hpf (brain analysis), or 96 hpf (morphological analysis). (**B**-**D**) Lateral and dorsal views of 96 hpf larvae treated with 0.1% DMSO (**B**) or 0.2 μM (**C**) or 0.5 μM bixafen (**D**). (**E**) Scheme depicting the different morphological parameters measured: body length (black double-arrow), head size (purple area), eye diameter (red double-arrow) and eye-otolith distance (green double-arrow). (**F**-**I**) Body length (**F**), head size (**G**), eye diameter (**H**) and eye-otolith distance (**I**), of 96 hpf larvae exposed to 0.1% DMSO (n = 14), or 0.2 μM (n = 15) or 0.5 μM (n = 12) bixafen. (**J**-**O**) Dorsal views of the brain of 36 (**J**-**L**) or 66 hpf transgenic Tg[*huC:G/U:RFP*] embryos (**M-O**), treated with 0.1 % DMSO (**J**, **M**), or 0.2 (**K**, **N**) or 0.5 μM bixafen (**L**, **O**), and showing the microcephaly of embryos exposed to low/medium doses bixafen. (**P-Q**) Measurements of brain volume of 36 (**P**) or 66 hpf (**Q**) embryos exposed to 0.1% DMSO (n_36hpf_ = 8; n_66hpf_ = 5), or 0.2 μM (n_36hpf_ = 6; n _66hpf_ = 5) or 0.5 μM bixafen (n_36hpf_ = 8; n_66hpf_ = 4), further illustrate the microcephaly of embryos exposed to bixafen at 96 hpf LC30 and LOAEL concentrations. ****, *p* < 0.0001; ***, *p* < 0.001; **, *p* < 0.01; *, *p* < 0.05; n.s., non-significant. Scale bar: (**B**-**D**) = 0.5 mm; (**J**-**O**) = 100 μm. Abbreviations: Tel, telencephalon; OT, optic tectum; Cer, cerebellum; SC, spinal cord.

### 2. Brain defects in embryos exposed to bixafen

Because several investigations have reported that exposure to SDHIs may induce neurotoxicity (Wang et al. 2020; Yao et al. 2018), we took advantage of the Tg[*elavl3:Gal4/5xUAS:RFP*] double transgenic line (Akerboom et al. 2012; Asakawa et al. 2008), hereafter referred to as Tg[*HuC:G/U:RFP*], a line in which all post-mitotic brain neurons are labelled by the fluorescent RFP protein, to precisely examine the morphology of the brain of 36 and 66 hpf embryos, following exposure to 0.2 and 0.5 μM bixafen. Live imaging of the brain of 36 and 66 hpf embryos, clearly indicated that exposure to both 0.2 and 0.5 μM bixafen, induced smaller brain size (Figure 1J-O). Precise quantification of brain volumes (Imaris software, Bitplane), indicated that 36 hpf embryos exposed to 0.2 (3.07 ± 0.13 x 10⁶ μm³) or 0.5 μM bixafen (1.60 ± 0.13 × 10^6^ μm^3^), displayed a significantly smaller brain compared with that seen in age-matched controls (3.57 ± 0.07 × 10^6^ μm^3^) (Figure 1P). Similarly, 66 hpf embryos exposed to 0.2 (7.45 ± 0.24 × 10^6^ μm^3^) or 0.5 μM bixafen (3.64 ± 0.15 × 10^6^ μm^3^), also displayed small brains compared to age-matched controls (8.51 ± 0.25 × 10^6^ μm^3^) (Figure 1Q). Our data thus showed that exposure of zebrafish embryos to low-dose bixafen corresponding to LOAEL induced microcephaly, a phenotype exacerbated in individuals incubated in 0.5 μM bixafen.

### 3. Impaired mobility and spinal motor neuron axon outgrowth defects in embryos exposed to bixafen

The small size of the brain observed in embryos exposed to bixafen suggests a neurotoxic effect of this SDHI. To further investigate this issue and precisely assess the consequences of bixafen exposure on brain functioning, we studied and compared the locomotor activity of 0.2 and 0.5 μM bixafen-exposed and control embryos by measuring the distance swum by these individuals over a 25-min period, using a ZebraBox tracking automaton (ViewPoint^®^). Results indicated that embryos treated with 0.2 or 0.5 μM bixafen, displayed reduced mobility, as shown by the distance traveled over a 25 minute period 77.3 ± 5.7 mm (N = 4, n = 72) and 52.7 ± 4.4 mm (N = 4, n = 77), respectively, compared with the distance travelled by controls (97.5 ± 4.9 mm, N = 4, n = 85) (Figure 2B-C). Interestingly, the hypoactive behavior of embryos exposed to 0.2 μM bixafen was markedly worsened in those exposed to 0.5 μM, which were almost motionless.

**Figure 2:**
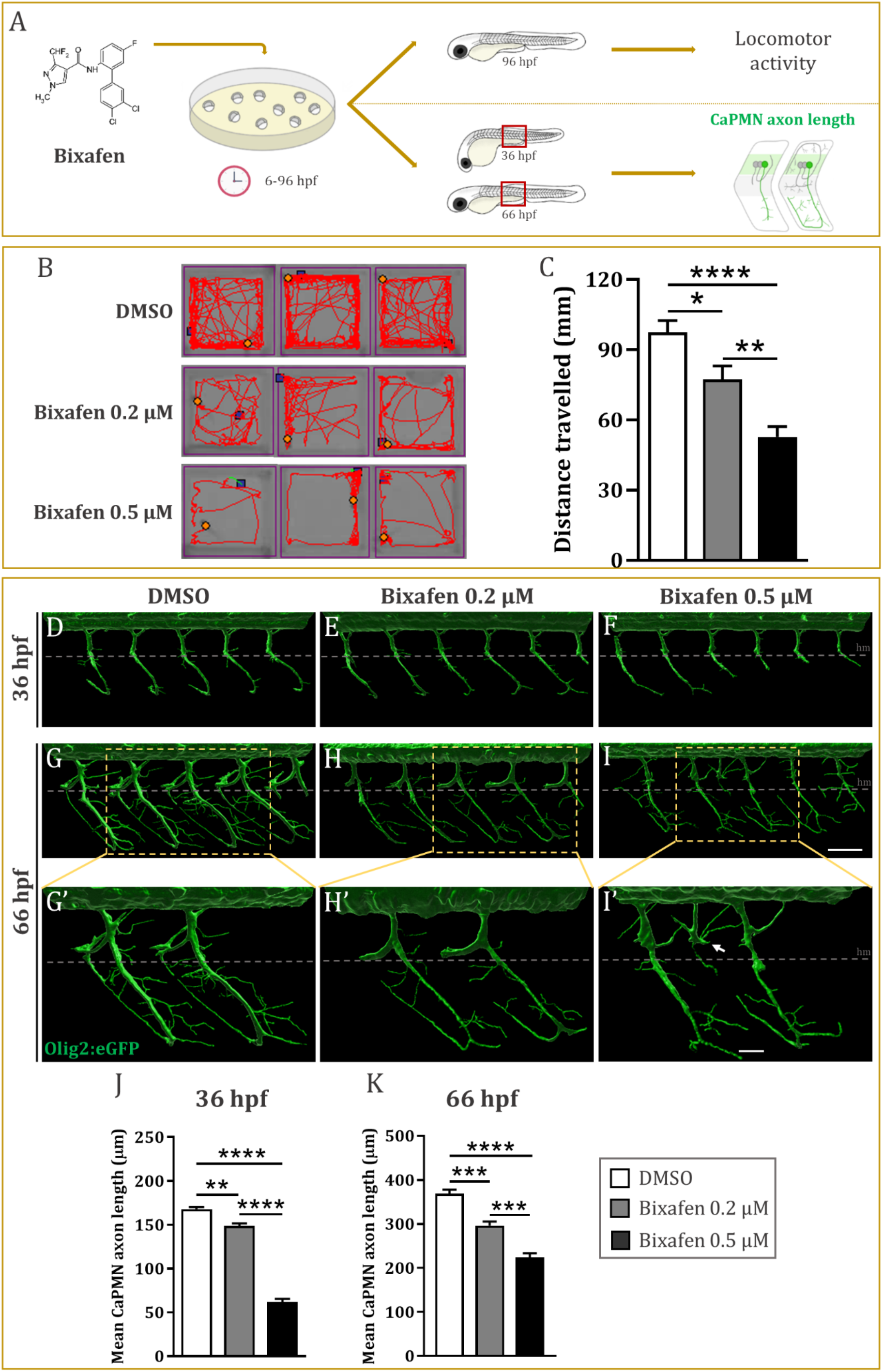
Bixafen exposure induces motor deficit and motor neuron axon defects. (**A**) Experimental setup used to analyze the motor activity and patterning of caudal primary motor neuron (CaPMN) axons and branches, in zebrafish embryos exposed to 0.2 or 0.5 μM bixafen, from 6 hpf onward up to 36 or 66 hpf (analysis of CaPMN axon morphology), or 96 hpf (analysis of motor activity). (**B**) Tracking plots illustrating the distance traveled by 96 hpf larvae treated with 0.1% DMSO (n = 85), or 0.2 (n = 72) or 0.5 μM bixafen (n = 77), during a 25 min time period. (**C**) Quantification of the distance traveled by 96 hpf larvae exposed to 0.1% DMSO, or 0.2 or 0.5 μM bixafen, during a 25 min time period. (**D**-**I**) Three-dimensional reconstruction of the organization of CaPMN axons and branches in 36 (**D**-**F**) or 66 hpf transgenic Tg[Olig2:eGFP] embryos (**G**-**I’),**exposed to 0.1% DMSO (**D**, **G**), 0.2 (**E**, **H**) or 0.5 μM bixafen (**F**, **I**), showing the disorganization of CaPMN axons and branches, following exposure to 0.2 and, even more, 0.5 μM bixafen (white arrow). (**G’**-**I’**) Higher magnifications of the morphology of CaPMN axons and branches, in embryos exposed to 0.1% DMSO (**G’**), or 0.2 (**H’**) or 0.5 μM bixafen (**I’**). (**J**, **K**) Quantification of the length of the primary CaPMN axons in 36 (**J**) and 66 hpf embryos (**K**), treated with DMSO (n_36hpf_ = 13; n_66hpf_ = 7), or 0.2 (n_36hpf_ = 8; n_66hpf_ = 6) or 0.5 μM bixafen (n_36hpf_= 22; n_66hpf_ = 9), showing significantly shorter axons following exposure to bixafen at LOAEL (0.2 μM) and, even more, 96hpf LC30 (0.5 μM). ****, *p* < 0.0001; ***, *p* < 0.001; **, *p* < 0.01; *, *p* < 0.05. Scale bar: (**D**-**I**) = 100 μm; (**G’**-**I’**) = 50 μm.

Because bixafen exposure induces brain defects, we next determined whether the behavioral deficits observed in bixafen-exposed embryos was a consequence of defects in spinal cord motor neurons or their axons. To address this issue, we next analyzed the morphology of the spinal cord in 36 and 66 hpf embryos exposed to 0.2 and 0.5 μM bixafen, using the Tg[*Olig2:eGFP*] transgenic line (Shin et al. 2003), in which all motor neuron progenitors and their axons are labelled by eGFP. In particular, we focused on caudal primary motor neurons (CaPMN) and their axons, which are the most accessible.

Live imaging indicated that in 36 hpf controls, primary motor neuron axons extended ventrally to contact target muscles, forming a highly regular pattern (Figure 2D). By contrast, motor neuron axons were markedly shortened in 36 hpf embryos treated with 0.2 bixafen (Figure 2E), a phenotype markedly exacerbated in age-matched embryos exposed to 0.5 μM, bixafen (Figure 2F). Three-dimensional reconstruction of CaPMN and their axons in 66 hpf control embryos revealed a highly stereotyped pattern with axons extending ventrally and branches projecting dorsally and laterally along the dorsal myoseptum and forming a regular pattern in the space between the notochord and the myotome (Figure 2G, G’). By contrast, following exposure to 0.2 μM bixafen, 66 hpf embryos displayed significantly shorter axons (Figure 2H, H’), a phenotype significantly worsened in embryos exposed to 0.5 μM bixafen (Figure 2I, I’), which also showed a marked disorganization of motor neuron axons and their branches, which extended in all directions.

Measurements of the length of CaPMN axons confirmed that bixafen exposure induced a marked shortening of these axons. Whereas in control embryos, the length of CaPMN axons was 167.9 ± 2.4 μm and 368.3 ± 9.3 μm, at 36 and 66 hpf, respectively, in embryos exposed to 0.2 μM, these values were reduced to 148.6 ± 3.1 μm at 36 hpf and 296.4 ± 9.0 μm at 66 hpf. Moreover, these defects were exacerbated in embryos exposed to 0.5 μM bixafen, with a mean CaPMN axon length of 61.8 ± 3.7 μm at 36 hpf and 223.7 ± 9.6 μm at 66 hpf (Figure 2J, K). Thus bixafen is neurotoxic and impairs the outgrowth and patterning of spinal motor axons and their branches.

## Discussion

It is well-known that the developing CNS in vertebrates is especially sensitive and vulnerable to toxic chemicals, including phytosanitary agents, which are increasingly present in the environment. More specifically, in recent years, bixafen, a latest-generation succinate dehydrogenase inhibitor (SDHI), has become one of the most widely-used fungicides worldwide. However, the toxicity of bixafen has been under-appraised *in vivo* in vertebrate models so far, and especially its neurotoxicity for the developing CNS, an especially serious issue because of the complete lack of species-specificity of this latest-generation SDHI (Bénit et al. 2019). In the present work, we used zebrafish embryos as an *in vivo* model and demonstrate that bixafen exposure during embryogenesis induces a series of CNS defects.

In close agreement with previous investigations (Li et al. 2020; Wu et al. 2018), we first observed that zebrafish embryos exposed to bixafen showed clear developmental delays, as shown by morphological examination of embryos exposed to 0.5 μM bixafen and above, but also by a decrease in the eye-otolith distance and hypopigmentation, both observed in embryos incubated in 0.2 and 0.5 μM bixafen, corresponding to LOAEL and 96hpf LC30, respectively. Moreover, at the brain level, we observed that these two same concentrations induced brain defects and microcephaly, an observation in close agreement with the markedly reduced expression of the neuronal *neuroD* gene observed in zebrafish embryos treated with bixafen (Li et al. 2020). Microcephaly has been repeatedly observed in embryos exposed to many pesticides, including Maxim^®^ XL (Svartz, Meijide, and Pérez Coll 2016) and glyphosate (Paganelli et al. 2010), emphasizing the high vulnerability of the brain to these chemicals, and our study shows that SDHI bixafen also induces microcephaly, even at low doses.

In addition, using an automaton that quantifies the speed and distance traveled by embryos, we also observed that embryos exposed to 0.2 and 0.5 M bixafen displayed reduced motility and almost complete paralysis, respectively, at 96 hpf. The quantification of swimming behavior clearly showed motor deficits in embryos exposed to 0.2 μM of bixafen which worsens in embryos exposed to 0.5 μM of bixafen (Figure 2C), that likely results of neuron abnormalities, not developmental delay. As it is known that the motor neuron is a major cell type regulating swimming behavior in zebrafish during early life (Brustein et al. 2003), the motor defects seen in bixafen-exposed embryos prompted us to investigate the differentiation of motor neuron axons and branches in embryos exposed to 0.2 and μM bixafen. In control embryos, motor neurons extend their axons in a highly stereotyped manner during development. In particular, caudal primary motor neuron (CaPMN) axons exit the spinal cord and extend toward the ventral muscle. Interestingly, in embryos exposed to bixafen, we observed a reduction in axon outgrowth and defects in their branches, even in those exposed to 0.2 μM bixafen corresponding to LOAEL, indicating that the development of motor neuron axons is highly sensitive to bixafen. Although the toxic effects of bixafen on the CNS are probably due at least in part to the developmental delay, the disorganization of motor neuron axons and branches clearly reveals a neurotoxicity of bixafen in zebrafish embryos.

Bixafen is a methyl-pyrazole, one of the SDHI fungicides used to manage plant diseases and inhibit the respiration of pathogenic fungi by blocking succinate complex II in the mitochondrial respiratory chain. SDH is an enzyme involved in both oxidative phosphorylation and the tricarboxylic acid cycle, two processes that generate energy. This enzyme has been shown to be irreplaceable in mitochondrial and cell metabolism and also to be highly conserved among all fungi, plants and animal species (Rehfus et al. 2016; Veloukas, Markoglou, and Karaoglanidis 2013; Yamashita and Fraaije 2018). Any adverse change or inhibition of SDH activity can therefore lead to many diseases, including those that have been linked to mitochondrial dysfunction (Birch-Machin et al. 2000; Bourgeron 2015; Van Coster et al. 2003; Finsterer 2008; Horváth et al. 2006; Parfait et al. 2000).

Several epidemiological studies published in the last two decades suggest harmful effects of pesticides on human health (Merhi et al. 2007; Weichenthal, Moase, and Chan 2010). Pesticide poisoning is a serious health problem that disproportionately affects infants and children (Rauh et al. 2012). Pesticides are known to cause millions of acute poisoning cases per year (Bertolote et al. 2006). Human exposure to pesticides can occur environmentally, through consumption in food and water (van den Berg et al. 2012). Bixafen is considered to be highly persistent in the environment (EFSA 2012). The half-lives of bixafen were 105 days in loam, 316 days in sandy loam, 1235 days in slit loam, and 82 days in water (EFSA 2012). A number of studies show that prenatal and early childhood exposure to some pesticides is associated with neurodevelopmental and cognitive disorders (Muñoz-Quezada et al. 2013; Ross et al. 2013). Aside from SDH inhibition, very little is known about the consequences of SDHI exposure on human health, and it is not known how bixafen affects the development of the CNS. As it is the case for other SDHIs, the toxic effect of bixafen observed in this study may be linked to the basic disruption of mitochondrial respiration chain complex II activity and subsequent accumulation of reactive oxygen species and oxidative DNA damage. Results showed that a high concentration of bixafen induces reactive oxygen species production and subsequent DNA damage in human cell lines (Graillot et al. 2012). However, a recent study also demonstrated that bixafen exposure did not cause oxidative stress in zebrafish (Li et al. 2020), suggesting that induction of oxidative stress may not be the main mechanism underlying bixafen-induced neurotoxicity in zebrafish embryos. Thus the brain and spinal cord defects observed in embryos exposed to low-medium bixafen concentrations may be associated with the metabolic abnormalities reported in SDHI-exposed animals (Graillot et al. 2012; Qian et al. 2018; Wu et al. 2018; Yang et al. 2018).

In summary, our study provides new evidence of bixafen toxicity on neurodevelopment in a vertebrate model, and may help us take appropriate precautions to ensure protection against the neurotoxic effects of these substances.

## Materials and Methods

### 1. Ethics statement

All the animal experiments described in the present study were conducted at the French National Institute of Health and Medical Research (INSERM) UMR 1141 in Paris in accordance with European Union guidelines for the handling of laboratory animals (http://ec.europa.eu/environment/chemicals/lab_animals/home_en.htm). They were approved by the Direction Départementale de la Protection des Populations de Paris and the French Animal Ethics Committee under reference No. 2012-15/676-0069

### 2. Zebrafish lines and maintenance

Zebrafish were maintained at 26.5 °C in 14 h light and 10 h dark cycles. Embryos were collected by natural spawning, and 0.003% 1-phenyl-2-thiourea (PTU) was added at 1 dpf (day post-fertilization) for embryos which were imaged *in vivo* to avoid pigmentation. The following transgenic lines were used: Tg[*elavl3:Gal4*]^*zf349*^ (Akerboom et al. 2012), T[*5xUAS:RFP*]^*nkuasrfp1a*^ (Asakawa et al. 2008), Tg[*Olig2:eGFP*] (Shin et al. 2003).

### 3. Chemicals

Dimethyl sulfoxide (DMSO) hybrid-max sterile (Sigma, D2650) was diluted in fish water to a final concentration of 0.1% (v/v). Bixafen, (*N*-(3’,4’-dichloro-5-fluorobiphenyl-2-yl)-3-(difluoromethyl)-1-methylpyrazole-4-carboxamide) (Bixafen, (Sigma-Aldrich, St. Louis, MO, USA, code: 32581, CAS-No: 581809-46-3, Lot# BCBS9732V, purity > 98%).) was dissolved at a stock concentration of 400 mM in pure DMSO (100%) and conserved at −20 °C in aliquots until use.

### 4. Toxicity assay

Wild-type (AB line) and transgenic lines, Tg[*HuC:G/U:RFP*] or Tg[*Olig2:eGFP*] embryos were collected after spawning and washed twice with clean fish water. Fertilized and normal 6 hpf embryos were selected, distributed randomly in 6-well microplates, 30 embryos per well, following OECD TG 236 fish acute toxicity test (OECD 2013). Each well was filled with 6 mL of solution for each condition, with 12 conditions in all: fish water, 0.1% DMSO, 0.1 μM, 0.2 μM, 0.3 μM, 0.5 μM, 1 μM, 1.5 μM, 2 μM, 2.5 μM, 3 μM and 4 μM bixafen in 0.1% DMSO. Each day, dead embryos were selected under OECD guidelines (coagulation, no heartbeat or no somite) and discarded.

### 5. Locomotor activity

96 hpf larvae were distributed individually in a 96 well plate in 200 μl of fish water. The plate was then placed in the recording chamber of the zebrabox system for 30 min habituation before 25 min recording of larvae locomotion (Brenet et al. 2019). The results indicate the distance swum by the larvae during the recording.

### 6. Embryo imaging

36 hpf or 66 hpf embryos were anesthetized with 112 μg/mL 3-aminobenzoic acid ethyl ester (tricaine, Sigma), immobilized in 1% low melting-point agarose in the center of a 35 mm glass-bottomed dish (Corning), and covered with fish water containing 112 μg/mL tricaine. For embryo morphology analysis, bright field images were captured using a stereomicroscope (Zeiss). Live imaging of transgenic lines Tg[*HuC:G/U:RFP*] and Tg[*Olig2:eGFP*] was done using a Leica SP8 confocal scanning laser microscope equipped with a Leica 20x/0.75 multi-immersion objective.

### 7. Image analysis

Body length, head size, eye diameter and eye-otolith distance measurements from bright field embryo images were made using the ruler tool in ImageJ. Brain volume and spinal neuron CaPMN axon length three-dimensional reconstructions and quantifications were analyzed using Imaris Measurement Pro (Bitplane Inc.).

### 8. Statistics

All statistics were obtained on Prism5 (GraphPad) and assessed using a one-way ANOVA test followed by a Tukey post-test. All data are represented as means ± SEM.

## Author Contributions

A.B. and R.H.A. performed the experiments and the analysis, designed the figures, and wrote the original draft. N.S.Y. supervised the project and wrote the manuscript.

## Acknowledgments

We thank Christiane Romain (Inserm UMR 1141), Olivier Bar (Inserm UMR 1141) for their technical assistance.

