## Supplementary data for "Bixafen, a succinate dehydrogenase inhibitor fungicide, causes microcephaly and motor neuron axon defects during development"

**Running title:** Neurodevelopmental defects in bixafen-exposed embryos

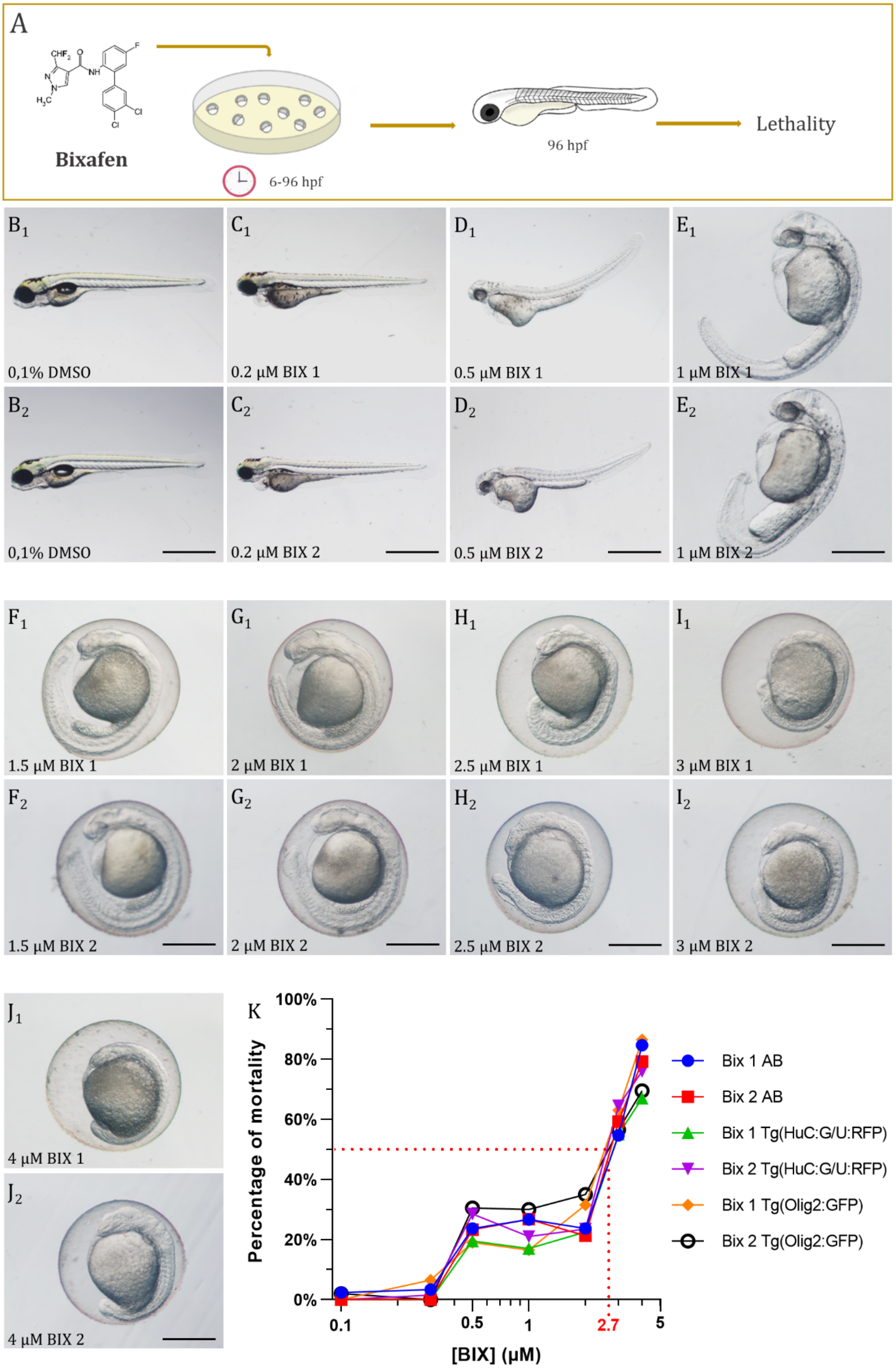
A
Bixafen
6-96 hpf
96 hpf
Lethality
B1
0,1% DMSO
C1
0.2 µM BIX 1
D1
0.5 µM BIX 1
E1
1 µM BIX 1
B2
0,1% DMSO
C2
0.2 µM BIX 2
D2
0.5 µM BIX 2
E2
1 µM BIX 2
F1
1.5 µM BIX 1
G1
2 µM BIX 1
H1
2.5 µM BIX 1
I1
3 µM BIX 1
F2
1.5 µM BIX 2
G2
2 µM BIX 2
H2
2.5 µM BIX 2
I2
3 µM BIX 2
J1
4 µM BIX 1
J2
4 µM BIX 2
K
Percentage of mortality
100%
80%
60%
40%
20%
0%
0.1
0.5
1
2.7
5
[BIX] (µM)
Bix 1 AB
Bix 2 AB
Bix 1 Tg(HuC:G/U:RFP)
Bix 2 Tg(HuC:G/U:RFP)
Bix 1 Tg(Olig2:GFP)
Bix 2 Tg(Olig2:GFP)

**Supplementary Figure 1: Bixafen toxicity**

(**A**) Experimental setup used to analyze bixafen toxicity, with concentrations of bixafen ranging from 0.2 to 4 µM, and embryo incubation from 6 hpf onward up to 96 hpf. (**B**-**J**) Morphology of 96 hpf larvae following exposure to two distinct batches of 0.1% DMSO (**B1-2**), or 0.2 ($\mathbf{B_1}$, $\mathbf{B_2}$), 0.5 ($\mathbf{D_1}$, $\mathbf{D_2}$), 1 ($\mathbf{E_1}$, $\mathbf{E_2}$), 1.5 ($\mathbf{F_1}$, $\mathbf{F_2}$), 2 ($\mathbf{G_1}$, $\mathbf{G_2}$), 2.5 ($\mathbf{H_1}$, $\mathbf{H_2}$), 3 ($\mathbf{I_1}$, $\mathbf{I_2}$) or 4 µM bixafen ($\mathbf{J_1}$, $\mathbf{J_2}$). (**K**) Curves of lethality of 96 hpf embryos from the wild-type AB and transgenic Tg[HuC:G/U:RFP] and Tg[Olig2:eGFP] lines, following incubation in increasing concentrations of bixafen ranging from 0.1 to 4 µM. The deducted 96 hpf LC50 of bixafen is 2.7 µM as indicated in red. Scale bar: (**B**-**D**) = 1 mm; (**E**-**J**) = 0.5 mm.
